# Shadows over Caribbean reefs: Identification of a new invasive soft coral species, *Xenia umbellata*, in southwest Puerto Rico

**DOI:** 10.1101/2024.05.07.592775

**Authors:** Daniel A. Toledo-Rodriguez, Alex Veglia, Nilda M. Jimenez Marrero, Joyce M. Gomez-Samot, Catherine S. McFadden, Ernesto Weil, Nikolaos V. Schizas

## Abstract

In October 2023, several colonies of an alien soft coral species were reported on shallow reefs in southwest Puerto Rico. The soft coral was identified as a xeniid octocoral (species undetermined), resembling the octocoral *Unomia stolonifera*, which has invaded and overgrown reefs in Venezuela in recent years. To conclusively characterize the species of the invading xeniid, we employed multilocus barcoding targeting four genes (ND2, mtMutS, COI, and 28S) of three separate colonies across three locations in southwest Puerto Rico. Sequence comparisons with xeniid sequences from GenBank, including those from the genera *Xenia* and *Unomia*, indicated a 100% sequence identity (>3,000 bp combined) with the species *Xenia umbellata* (Octocorallia : Malacalcyonacea : Xeniidae). *Xenia umbellata* is native to the Red Sea and to our knowledge, this represents the first confirmed case of this species as an invader on Caribbean reefs. Similar to *U. stolonifera, X. umbellata* is well known for its ability to rapidly overgrow substrate as well as tolerate environmental extremes. In addition, *X. umbellata* has recently been proposed as a model system for tissue regeneration having the ability to regenerate completely from a single tentacle. These characteristics greatly amplify *X. umbellata’*s potential to adversely affect any reef it invades. Our findings necessitate continued collaborative action between local management agencies and stakeholders in Puerto Rico, as well as neighboring islands, to monitor and control this invasion prior to significant ecological perturbation.

## Introduction

Globally, reports of invasive marine species have increased, posing significant threats to native marine organisms and the ecosystems they inhabit (Bailey, 2015). Invasive organisms transform their new host habitats, often inducing structural and functional shifts within ecosystems, resulting in substantial ecological and economic repercussions (Alidoost Salimi et al., 2021; Menezes et al., 2021; Molnar et al., 2008). Lacking natural predators in the invaded regions, these species may alter vital local food webs and cause trophic shifts, exacerbating already ongoing challenges for marine biodiversity and resource conservation and management (Menezes et al., 2021; Molnar et al., 2008; Simberloff & Von Holle, 1999; Simkanin et al., 2012; Williams & Grosholz, 2008; Zhan et al., 2015). The economic repercussions caused by invasive species are significant as well, often being introduced to productive, yet vulnerable habitats like coral reefs which are essential for coastal protection, tourism, and fishing (Groeneveld et al., 2018; Pavliska, 2019; Pimentel et al., 2001; Simkanin et al., 2012; Zhan et al., 2015). Coral reef invasives, like the infamous lionfish (*Pterois volitans*) now a fixture on Caribbean reefs, are introduced through human activities such as shipping and irresponsible practices in the aquarium trade (Mantellato et al., 2018; Menezes et al., 2022; Minchin, 2006). Because invasive species pose such a serious threat, investigating suspected invasions and accurately identifying the culprit species is crucial for strategic intervention to prevent the use of unsuitable tactics and protect these important resources (Anderson et al., 2015; Groeneveld et al., 2018; Molnar et al., 2008; Pavliska, 2019; Ruiz-Allais et al., 2021; Zhan et al., 2015).

During recent years, Caribbean reefs have dealt with several invasions of an octocoral species identified as *Unomia stolonifera* (Family Xeinidae; Gohar, 1938), a species native to the Indo-Pacific region. In Venezuela, from Coronas Bay to Arapo Island, *U. stolonifera* has become a major problem, spreading rapidly and becoming a dominant organism on local reefs displacing the native sessile fauna. *Unomia stolonifera* is characterized by its biofouling capacity; it has occupied between 30% and 80% of the benthic habitats it has invaded, reducing local biodiversity, especially coral cover, thus demonstrating the urgent challenges caused by marine invasions (Benayahu et al., 2021; Ruiz-Allais et al., 2014; Ruiz-Allais et al., 2021). In Cuba, a pulse coral presumed to be *U. stolonifera* was reported on reefs in 2022, and efforts to control and eradicate the invader was successful in some localized areas demonstrating that this invasion can be impeded (Espinosa et al., 2023). Although, the battle is still ongoing in other coastal regions around the island, highlighting the persistence of these pulse corals. Therefore, it is imperative to catch a pulse coral invasion early to provide the best chance for successful mitigation. Recently in October 2023, an unknown xeniid species was observed on reefs in south Puerto Rico, raising significant concerns about the possible presence of the notorious *U. stolonifera* in the region.

Since the initial sighting, the Puerto Rico Department of Natural and Environmental Resources (PRDNER), through a public awareness campaign, was able to corroborate additional patches of pulsing xeniids on reefs in the south and southwest Puerto Rico. Colonies of the unidentified octocoral have so far been observed on reefs near Ponce, Guayanilla, Guánica and within the La Parguera Natural Reserve (LPNR). The invasive octocoral colonies were found growing on various substrata, often smothering native zoanthids, corals and sponges (Fig. 1). Given the aggressive nature and biofouling capabilities of previously reported invasive xeniid species (Mantellato et al., 2018; Menezes et al., 2022; Ruiz Allais et al., 2021), it is crucial to characterize the exotic species and understand its potential impact as an invader. This study aims to provide a definitive taxonomic identification of the invading Xeniidae species using a multilocus barcoding approach, thereby contributing essential information for developing effective monitoring and mitigation strategies. In addition to clarifying the species identity, this work seeks to communicate its potential as an invader on Caribbean reefs, providing a foundation for informed management and policy decisions in affected regions.

**Figure 1:**
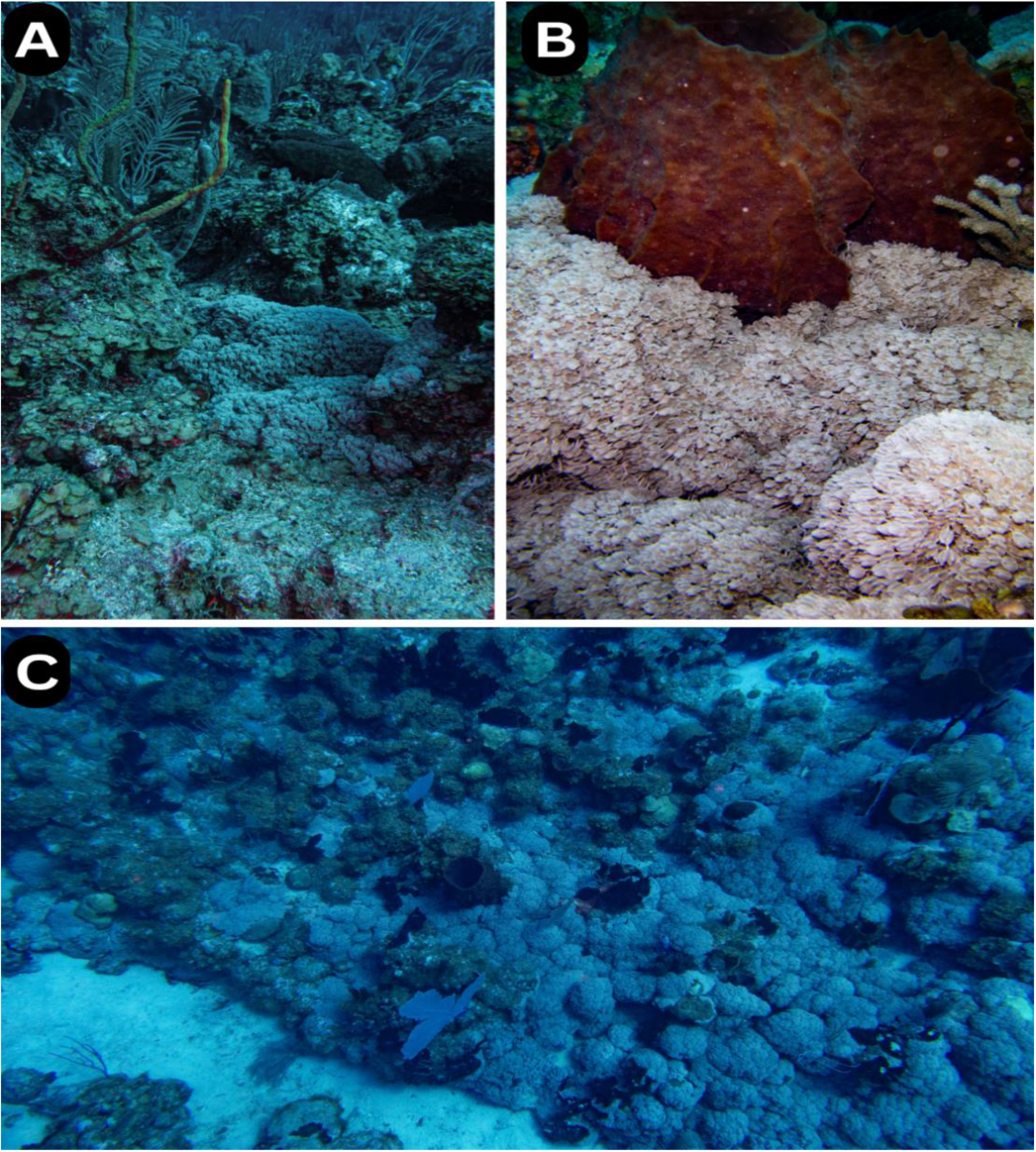
*Xenia umbellata* overgrowing the reefs. Image A: shows *X. umbellata* colonizing both bare substrate and areas previously occupied by macroalgae. B: captures the early stages of *X. umbellata* overgrowing a barrel sponge (*Xestospongia muta*). C: Over the top view of *X. umbellata* expanding over the reef. Photos by Daniel A. Toledo-Rodriguez.

## Methods

### Study Sites

Three sample locations were selected based on the reported presence of the xeniid soft coral along the southern coast of Puerto Rico. Colonies were sampled from sites located in La Parguera Natural Reserve (LPNR) (n=2) and Ponce (n=1) (Table 1). The LPNR sample sites (n=2) are deep spur and groove reefs, ranging in depths from 20-23 meters. The Ponce site, however, is a shallow reef on the northeast side of Caja de Muertos island. In both areas, the rapid spread of invasive octocoral has changed the surrounding environment with patches that have reduced the available substrate by overgrowing the reef (Fig. 1). The water current predominately flows westward along the south coast of Puerto Rico and the two sites in the LPNR are ∼47 km west from the colony reported and sampled in Ponce, which is near the commercial Port of Ponce (Table 1 and Fig. 2). All collections were conducted by PRDNER staff who had the authority to intervene with the invasive species and prioritized the genetic study among the response strategy activities.

**Table 1.** Sampling sites with sample ID, average depths, coordinates and distance between each sampling site.

| *Site* | **Sample ID** | **Depth (m)** | **(Latitude, Longitude)** | **Distance to Site 2**<br>**(km)** | **Distance to Site 3 (km)** |
| --- | --- | --- | --- | --- | --- |
| *Site 1 (LPNR)* | XE4b | 22 | N 17.89582, W 66.9728 | ~0.08 | ~47 |
| *Site 2 (LPNR)* | Xe5a | 20 | N 17.89578, W 66.9736 | - | ~47 |
| *Site 3 (Ponce)* | XP2a | 2 | N 17.8951, W 66.5190 | ~47 | - |

**Figure 2:**
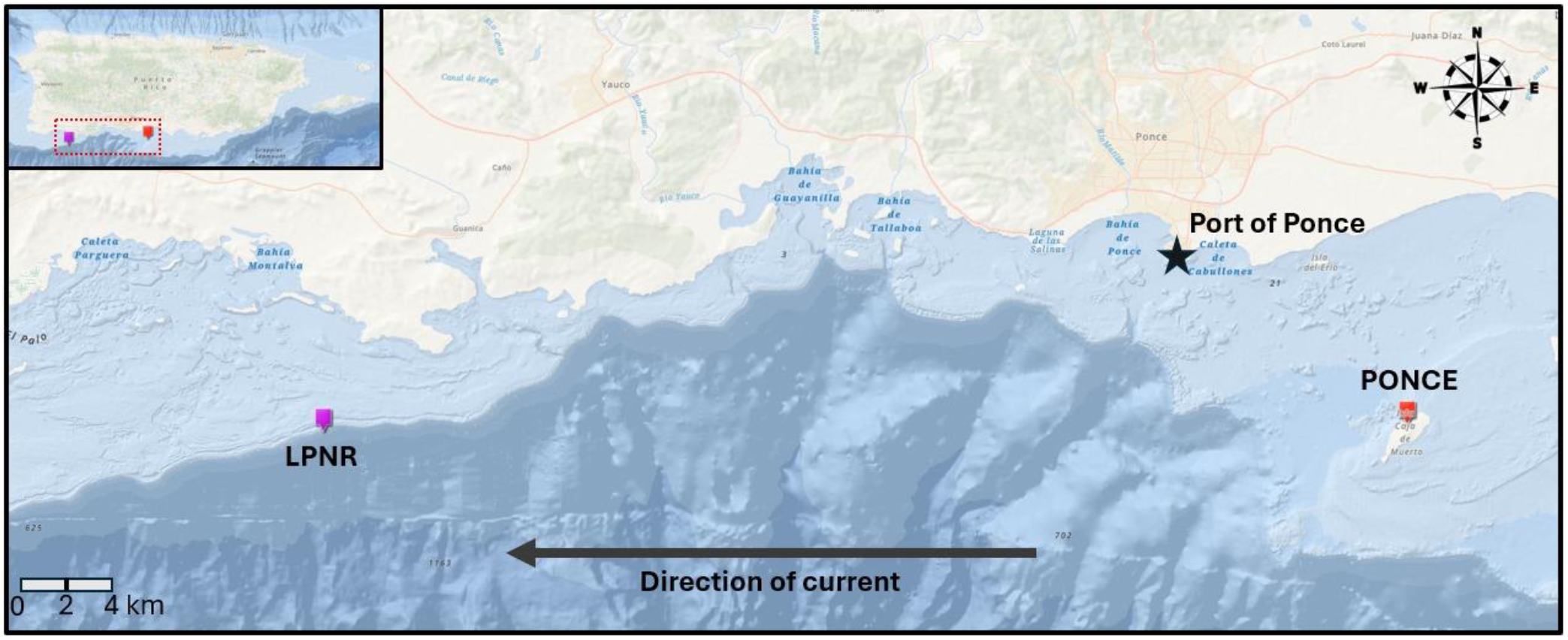
Map showing southwest Puerto Rico sampling region with the locations of the colonies sampled in the LPNR (purple) and Ponce (red). The black arrow depicts the direction of the water current in the area. This figure shows a visual representation of how distant both study sites are from each other (approximated distance in km shown in Table 1).

### Sample collection

All samples were collected from the sites provided in Table 1 (see Fig. 2 for visual reference), using SCUBA and/or snorkeling. Two samples were collected per colony (Fig. 3 and 4), carefully extracted either manually or with tweezers, and then placed in a 50mL tube, minimizing impacts on the surrounding area and preventing potential fragmentation which could promote the spread of the soft coral. Collection date, depth, substrate type, and additional observations were recorded on a diving slate. One fresh tissue sample (whole polyps) was stored in a -80ºC freezer and several other polyps were preserved in 95% ethanol as vouchers at the Marine Genomic Biodiversity Lab at Magueyes Island.

**Figure 3:**
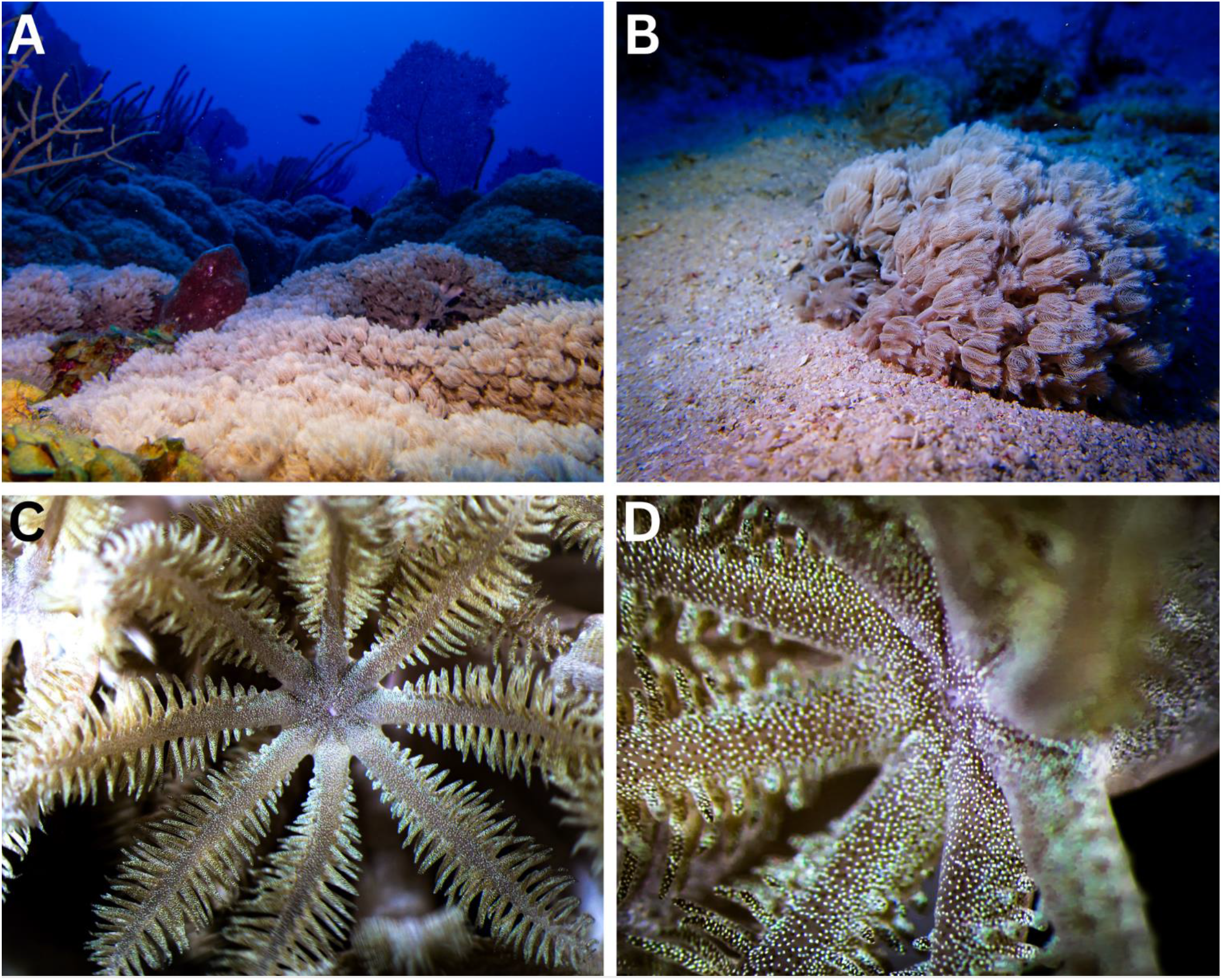
Images of the invasive octocoral *Xenia umbellata* overgrowing local coral reefs from LPNR. A: Site 1 showing *X. umbellata* dominance of the benthic landscape. B: A xeniid colony from Site 2 growing in coarse sand. C: A macro-picture of a xeniid polyp from Site 1, presenting the mouth and pinnate tentacles. D: a close-up image of a polyp observed in Site 2. Photos by Daniel A. Toledo-Rodriguez.

**Figure 4:**
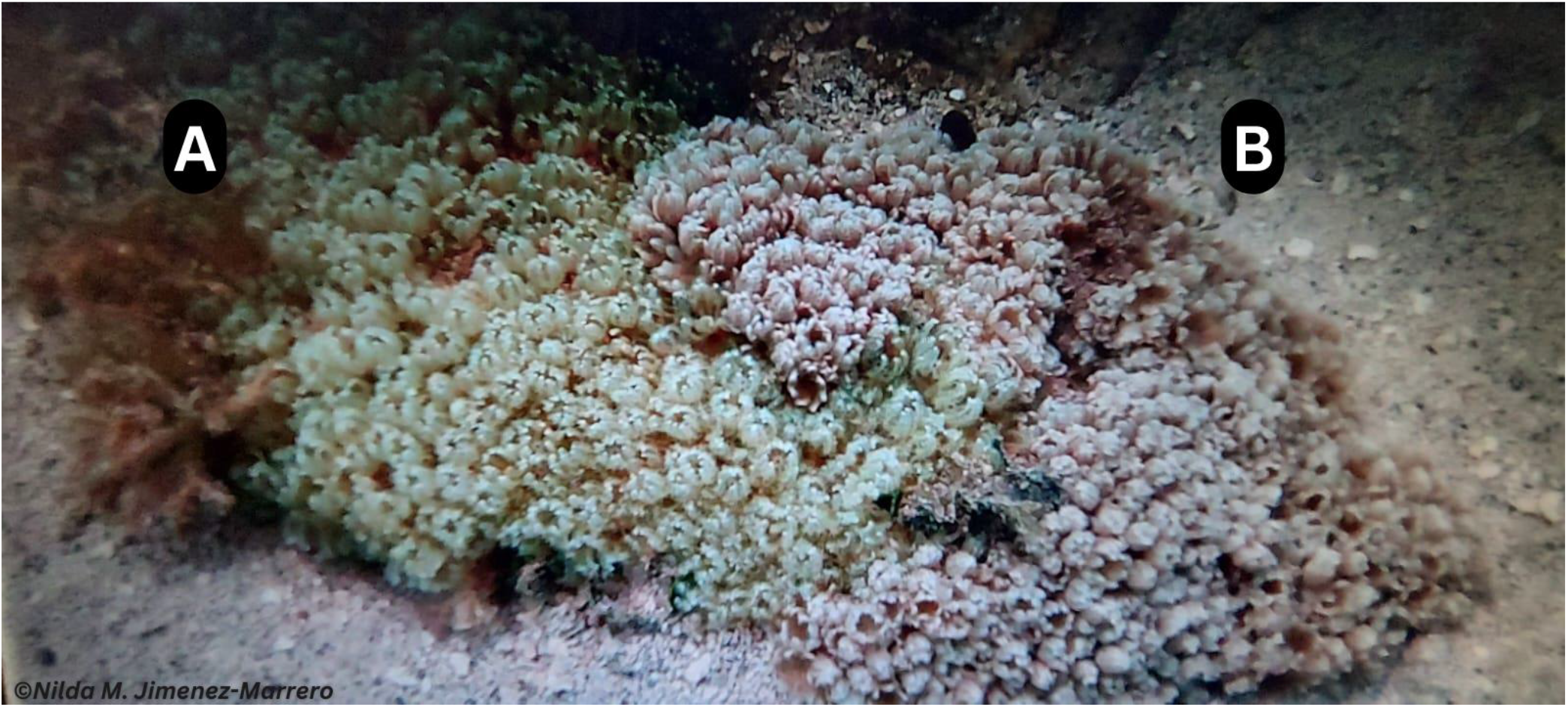
Two xeniid colonies from Site 3, located in Ponce, Puerto Rico, exhibiting color variation. The colony on the right side is genetically confirmed as *Xenia umbellata*. The species identification of the colony on the left side has not been confirmed yet but it is likely the same xeniid species. Photo provided by Dr. Nilda M. Jimenez-Marrero.

**Figure 5:**
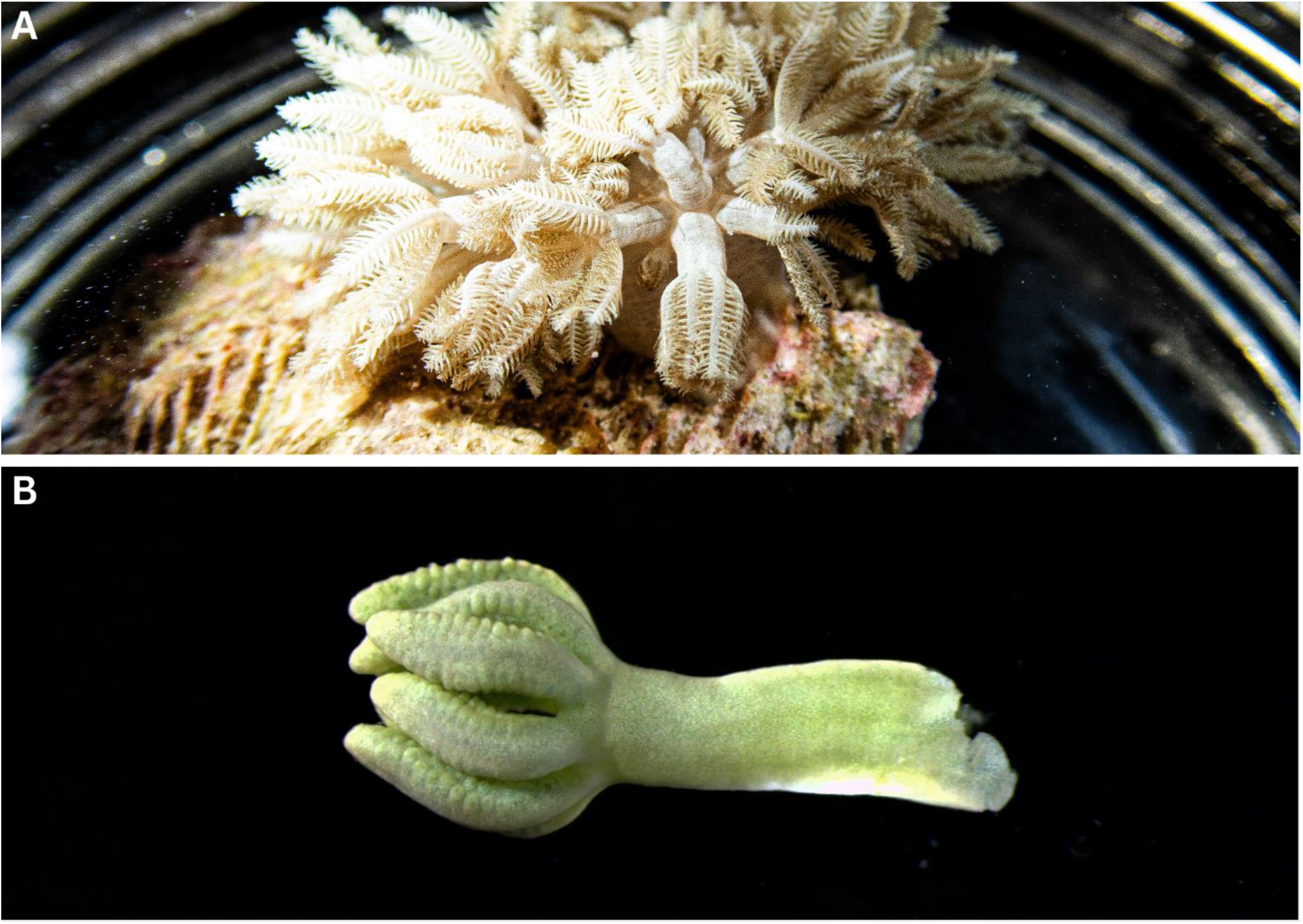
Photos of *Xenia umbellata* samples being processed under a dissecting microscope for DNA analysis. A: A live sample from Site 1 in a glass container submerged in ocean water. B: A close-up picture of one polyp submerged in 95% ethanol and observed under the microscope. Photos by Daniel A. Toledo-Rodriguez.

### DNA extraction, amplification, and sequencing

For a DNA template, 4-5 tentacles were excised from each xeniid polyp (Fig. 4) using sterile tweezers and a scalpel. DNA extraction was performed using the Qiagen DNeasy PowerSoil® kit according to the manufacturer’s instructions, with an additional homogenization step consisting of three sets of 20 seconds, with 1-minute intervals on ice between the sets. This process was employed to maximize DNA yield. The quantity and quality of DNA were checked using a NanoDrop 2000™ Spectrometer. PCR (Polymerase Chain Reaction) amplifications of three mitochondrial genes (16S/ND2, mtMutS, COI) and the nuclear large ribosomal subunit (28S rRNA) were done with the primers listed in Koido et al. (2022). The mixture of the PCR reactions with a total volume of 25µl contained 1.5µl DNA, 8.5µl ddH2O, 1.25µl of each primer (10µM), and 12.5µl KAPA Taq ReadyMix (Biosystems company). All reactions were conducted on a Bio-Rad T100™ Thermal Cycler. The PCR protocol for the three mitochondrial and the nuclear markers was 95°C for 3 minutes, followed by 35 cycles of 90 sec at 94°C, 60 sec at 58°C, and 60 sec at 72°C. Amplicons were verified on a 1.4% agarose gel electrophoresis stained with ethidium bromide and visualized in UV fluorescence. Sanger Sequencing Services were provided by the Sequencing and Genomics Facility of the University of Puerto Rico, Rio Piedras. Amplified DNA was sequenced in both directions in an Applied Biosystems SeqStudio Genetic Analyzer using the Big Dye 3.1 Terminator Cycle Sequencing Kit. The DNA traces were inspected for quality and accuracy in nucleotide base assignment in Codon Code Aligner v. 10.0.2 (Codon Code Corp.). Sequences were trimmed and then aligned with Clustal Omega (Sievers et al., 2011) as implemented in Codon Code Aligner. All DNA sequences have been submitted to GenBank (Accession Numbers XXXXX-XXXXX).

## Results and Discussion

Notorious for their invasive behaviors, xeniid species exemplify the adaptability and competitiveness in non-native environments (Ruiz-Allais et al., 2014; 2021; Studivan et al., 2015) that characterize many invasive species, disrupting local marine habitats and outcompeting local species (Benayahu et al., 2021; Mantellato et al., 2018; Menezes et al., 2022; Ruiz-Allais et al., 2021). Understandably, the detection of the xeniid colonies in southwest Puerto Rico generated stress on the local community, scientists and government agencies due to possible ecological impacts. Early detection and correct taxonomic identification of the invasive species was paramount to start learning about the biology of the species in its native range and make predictions about its ecological interactions with the benthic ecosystems of Puerto Rico. Utilizing a multilocus barcoding approach, we examined the genetic sequences of three colonies of the invasive xeniid octocoral to determine their species identity and assess potential genetic variation. The length of sequences after end trimming was reduced to 690bp (ND2), 844bp (COI), 715bp (mtMutS), and 773bp (28S). There was no genetic variation among the three examined xeniid colonies in >3,000bp of combined sequences. BLASTn searches of the Puerto Rican xeniid sequences for all four genes revealed 100% sequence identity (e.g., GenBank Accession Numbers, 16S/ND2: KY442428; COI: KC864981; mtMutS: PP470895; 28S: MK400153 - Haverkort-Yeh et al., 2013; McFadden et al., 2017, 2019) with the species *Xenia umbellata* (Lamarck, 1816). In addition, sequences of all four genes were compared against other published and unpublished xeniid sequences (McFadden unpub. data) verifying the initial species diagnosis. These findings reveal a new invasive octocoral on Caribbean reefs, and one that has the characteristics to incur significant long-term ecological impacts in affected localities and regions (Fig. 1C).

*Xenia umbellata* is endemic to the Red Sea (McFadden et al., 2019) and its presence in Puerto Rican waters warrants attention from management agencies and the public because of its ecological attributes. All climate change scenarios predict warmer, more acidic oceans exposing marine life rapidly to environments near or beyond their physiological limits. *Xenia umbellata* has shown significant resistance in lab experiments to warmer, more acidified conditions. In physiological experiments, *X. umbellata* exhibited resistance to warming waters (Thobor et al., 2022) which could potentially increase in eutrophic environments (Vollstedt et al., 2020). In the field, observation of appraently healthy *X. umbellata* colonies in October 2023 and in early 2024, during the most extensive and intensive thermal anomaly to impact the southwestern PR coastal habitats which produced significant hard coral mortalities (Weil unpublished data), further indicates *X. umbellata’s* resistance to high thermal stress. Furthermore, differential pulsation of tentacles (a charismatic attribute of the species that makes it desirable in the aquarium trade) may permit *X. umbellata* to tolerate a wide range of pH with few effects on its overall health (Tilstra et al., 2023). Taken together, the local field observations and the previously produced experimental data indicate that under ocean warming and ocean acidification scenarios, *X. umbellata* may have a competitive advantage over native corals and other reef organisms. The potential of this species to dominate the benthos is exemplified by its promotion as a novel model organism for studying regeneration in octocorals because of its ability to regenerate from single tentacles or polyps (Nadir et al., 2023). Divers in southwestern Puerto Rico have observed *X. umbellata* colonies growing on bare substrates such as rocks, rubble and sand (Fig 2B), over and under scleractinian corals (Fig. 3A) such as *Pseudodiploria strigosa* and *Colpophyllia natans*, around soft corals and sponges, over macroalgae (Fig. 1A) and inside crevices, confirming its potential to dominate the benthic landscape.

Common routes of transmission for invasive marine species include the aquarium trade, climatic events, ballast water, commercial shipping and boat hulls, and plastic or other floating debris (e.g., coconut husks, sargassum, plastic bottles) (Olden et al., 2021; Bailey 2015; Mantelatto et al., 2018; García-Gómez et al., 2021). *Xenia umbellata* is a common species in the aquarium trade because of its beautiful pulsating tentacles, the ability to regenerate from small tissue sections, and its hardiness during transportation. It is possible that, like lionfish, this species was released into local water ways, permitting the invasion. One report of *X. umbellata*, however, is near the Port of Ponce (Fig. 2), which services international commercial ships. This suggests that discharged ballast water or debris from a vessel hull is a plausible alternative route for long-distance transport of the soft coral to Puerto Rico. Several other non-native corals that have recently colonized the Caribbean appear to have been transported as fouling organisms on semi-submersible platforms (Hoeksema et al., 2023). Understanding the exact route of introduction is crucial for implementing effective monitoring and management strategies and should be a direction of future interrogation.

Personnel from the Department of Natural and Environmental Resources and HJR Reefscaping have initiated efforts to remove *X. umbellata* colonies to prevent the further spread of the soft coral. The PRDNER has been engaging and creating awareness so that marine scientists, stakeholders, and local communities are aware of the species and its potential to damage coastal ecosystems, urging all to report it if observed. Future work will include genetic work to confirm the species reported in Cuba providing key insight on propagation venues and promoting responsible practices within the aquarium trade and other activities that may facilitate spread (e.g., dredging, bottom-fishing gear, recreational boating, and anchoring). By fostering collaboration between scientists, local agencies, and the public, we can implement comprehensive monitoring and management strategies to safeguard Puerto Rico’s reefs and mitigate the ecological and economic repercussions of this invasion.

## Acknowledgments

We extend our gratitude to Dr. Nilda M. Jiménez-Marrero of the Department of Natural and Environmental Resources, Puerto Rico, for their encouragement and invaluable support in providing the sampling permit. We also thank Dr. Hector J. Ruiz-Torres from HJR Reefscaping for providing the boat and diving tanks for the sample collection. This publication was made possible with support from the Sequencing and Genomics Facility of the UPR Río Piedras & MSRC/UPR, funded by NIH/NIGMS-Award Number P20GM103475. Finally, the Department of Marine Sciences, UPRM for logistical support.

